# FGF/MAPK signaling sets the switching threshold of a bistable circuit controlling cell fate decisions in ES cells

**DOI:** 10.1101/015404

**Authors:** Christian Schröter, Pau Rué, Jonathan Peter Mackenzie, Alfonso Martinez Arias

## Abstract

Intracellular transcriptional regulators and extracellular signaling pathways together regulate the allocation of cell fates during development, but how their molecular activities are integrated to establish the correct proportions of cells with particular fates is not known. Here we study this question in the context of the decision between the epiblast (Epi) and the primitive endoderm (PrE) fate that occurs in the mammalian preimplantation embryo. Using an embryonic stem (ES) cell model, we discover two successive functions of FGF/MAPK signaling in this decision. First, the pathway needs to be inhibited to make the PrE-like gene expression program accessible for activation by GATA transcription factors in ES cells. In a second step, MAPK signaling levels determine the threshold concentration of GATA factors required for PrE-like differentiation, and thereby control the proportion of cells differentiating along this lineage. Our findings can be explained by a simple mutual repression circuit modulated by FGF/MAPK signaling. This may be a general network architecture to integrate the activity of signal transduction pathways and transcriptional regulators, and serve to balance proportions of cell fates in several contexts.

## Introduction

To ensure the faithful development of multicellular organisms, cell fate decisions in populations of undifferentiated cells have to be tightly balanced. It is now well established that transcriptional networks and extracellular signals together control these decisions, but how their interactions determine the proportions of cells differentiating along particular lineages is often not known. This question is of particular importance in one of the first cell fate decisions in the mammalian preimplantation embryo, where a small number of inner cell mass (ICM) cells have to reliably populate both the epiblast (Epi) lineage that will give rise to the embryo proper, as well as the primitive endoderm (PrE) lineage, which differentiates into tissues that function in patterning and nutrient supply of the embryo (Rossant and Tam, 2009). The factors underlying this fate decision have been studied in embryos and embryonic stem (ES) cells (Cho et al., 2012; Fujikura et al., 2002; Shimosato et al., 2007), clonal derivatives of ICM cells that are biased towards the Epi fate but harbor a latent PrE differentiation potential (Beddington and Robertson, 1989; Morgani et al., 2013). Mutant analysis indicates that a transcriptional network centered on the transcription factor Nanog marks and defines the Epi fate (Chambers et al., 2007; Frankenberg et al., 2011), while a network centered on GATA transcription factors underlies the PrE fate (Bessonnard et al., 2014; Schrode et al., 2014). Nanog and GATA6 are co-expressed in early ICM cells, but their expression patterns become mutually exclusive as cells commit to specific fates (Plusa et al., 2008), suggesting mutually repressive interactions between the two programs (Frankenberg et al., 2011; Schrode et al., 2014). Studies using genetic mutants and pharmacological inhibitors have furthermore shown that the FGF/MAPK signaling pathway promotes the PrE fate at the expense of the Epi fate in embryos (Kang et al., 2012; Nichols et al., 2009; Yamanaka et al., 2010), and that it is required for PrE-like differentiation of ES cells (Cho et al., 2012). How the activities of the transcriptional networks are integrated with the activity of the FGF/MAPK signaling pathway, and how these inputs together control the proportion of cells differentiating along either lineage has not been systematically investigated.

Recently, a mathematical model for the decision between the Epi and the PrE fate has been proposed (Bessonnard et al., 2014), in which Nanog and GATA repress each other, and reinforce their own expression through direct positive feedback. This defines a dynamic system with three stable states in which cells either express GATA6 or Nanog alone, or co-express the two markers. In this model, FGF/MAPK signaling both promotes GATA6 expression and inhibits Nanog expression, and differences in FGF/MAPK signaling between cells have been proposed to underlie fate choice from the co-expression state (Bessonnard et al., 2014). While this model is consistent with static phenotypes of wildtype embryos and genetic mutants, the gene expression dynamics proposed by this model have not directly been tested. It is also not clear whether all proposed links are required to explain the behavior of the genetic circuit underlying this cell fate decision, and which one of the two inputs into the system – signaling and transcription factor activity - bears most on the fate decision. Addressing these open questions requires quantitative modulation of the inputs into the genetic circuit regulating fate choice, and following its dynamics in single cells in real time. Here we achieve this by transiently expressing fluorescently tagged GATA factors in ES cells carrying live reporters for the Epi and the PrE fate. This allows us to recreate a state of co-expression of Epi and PrE determinants akin to the state of ICM cells in the embryo, and to follow the resolution of this state in real time. We find that cells rapidly exit the co-expression state towards one of two mutually exclusive states, i.e. the system is bistable. PrE-like differentiation occurs in cells exposed to GATA factor levels above a threshold, and the function FGF/MAPK signaling is to set this threshold dose. This provides a mechanism through which both transcription factor activity and signaling can tune the proportions of cells with specific fates. Recapitulating the dynamic behavior of the circuit *in silico* only requires mutual repression between the transcriptional networks underlying the Epi and the PrE fates without any positive feedback loops, and a single repressive input of MAPK signaling on the Epi-specific program. This data-based model for the Epiversus-PrE fate decision, much simpler than previously proposed models, will serve as a basis to guide further experimental and theoretical exploration of this crucial fate decision of mammalian embryogenesis. Furthermore, our finding that FGF/MAPK signaling can balance the proportions of alternative fates in cell populations by setting the response threshold of a regulatory network to a transcription factor input is a novel principle for this signaling pathway which might be relevant in developing tissues beyond the ICM.

## Results

### An ES-cell model system to investigate PrE-like fate choice in culture

To model in culture the transition from Nanog;GATA6 co-expression to mutually exclusive expression of Epi and PrE markers that characterizes the Epi-versus-PrE fate decision (Plusa et al., 2008), we used a doxycycline-inducible system to transiently express GATA6-FLAG in ES cells (Beard et al., 2006; Mulvey et al., 2015; Wamaitha et al., 2015)(Fig. 1A). Individual cells co-expressed inducible GATA6-FLAG and endogenous Nanog protein after a 6 h doxycycline pulse (Fig. 1B). 24 h after doxycycline removal, the cells had degraded the exogenous GATA6-FLAG, but a subset now stained positive for the endogenous PrE marker GATA4 (Fig. 1C). Virtually all GATA4-positive cells were negative for Nanog staining, suggesting that following GATA6/Nanog co-expression, ES cells transition to one of two mutually exclusive states, marked by the expression of Epi- and PrE markers, respectively. This is similar to the behavior of ICM cells, and suggests that a previously reported stable state of co-expression of Nanog and endogenous GATA factors (Bessonnard et al., 2014) is not accessible in our system.

**Fig. 1:**
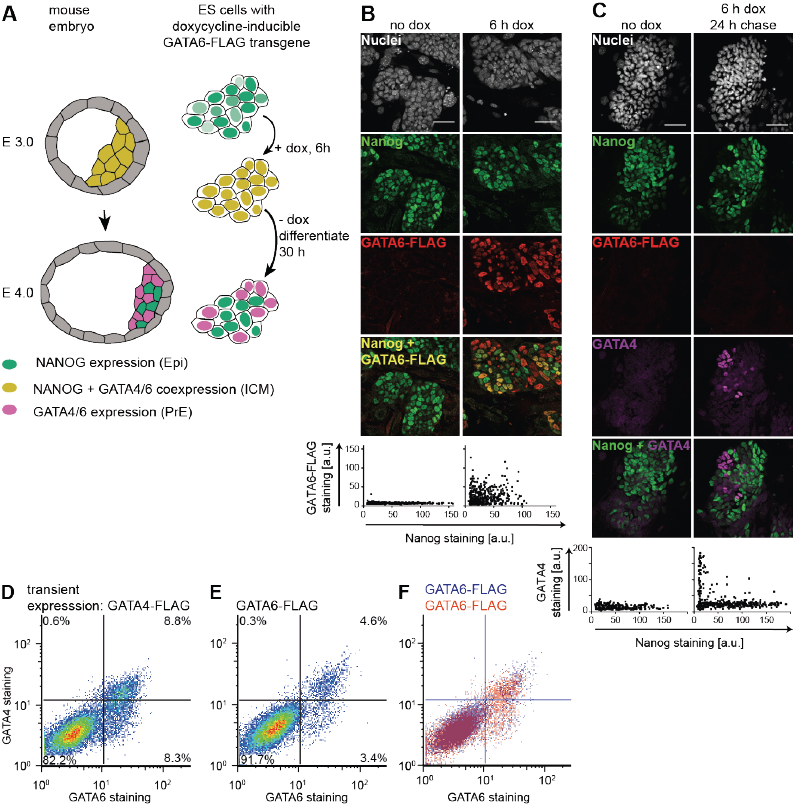
Expression of endogenous markers of PrE-like differentiation following transient expression of GATA6-FLAG and GATA4-FLAG. ***A*** *Experimental approach: Doxycycline-induced transgene expression creates a GATA6/Nanog co-expression state in ES cells similar to the situation in the ICM, from which cells can embark on PrE-like differentiation, or return to the Nanog-positive state.* ***B*** *Immunostaining (upper panel) and quantification (lower panel) of untreated (left) or doxycycline-treated (right) inducible ES cells indicates coexpression of Nanog and GATA6-FLAG in individual cells after 6 h of doxycycline treatment. Co-expression is limited because of heterogeneous Nanog and GATA6-FLAG expression in the presence of serum and feeders. Scale bar, 50 μm.* ***C*** *Immunostaining (upper panels) and quantification (lower panels) of Nanog and GATA4 expression 24 h after the end of a 6 h doxycycline pulse. GATA4 expression depends on doxycycline treatment, and is mutually exclusive with Nanog expression. Scale bar, 50 μm.* ***D, E, F*** *Flow cytometry of cells immunostained for endogenous GATA4 and GATA6 one day after transient GATA4-FLAG (****D****) or GATA6-FLAG (****E****) expression. F shows the overlay of* ***D*** *and* ***E***.

Consistent with previous studies (Fujikura et al., 2002; Mulvey et al., 2015; Shimosato et al., 2007), we found that transient expression of doxycycline-inducible GATA4-FLAG instead of GATA6-FLAG led to the same expression pattern of endogenous GATA factors, but doubled the proportion of differentiating cells (Fig. 1 D – F, supplementary material Fig. S1). This led us to induce PrE-like differentiation with GATA4 and to use endogenous GATA6 expression to monitor the differentiation event in all following experiments. Furthermore, we tagged the inducible GATA4 protein with an mCherry fluorescent protein. This did not compromise the activity of the fusion protein to induce PrE-like differentiation (supplementary material Fig. S2), and allowed us to follow the heterogeneous expression of the doxycycline-induced transgene in individual live cells.

### ES cell culture conditions affect the expression of endogenous PrE markers following a GATA4-mCherry pulse

For an induced transcription factor to trigger a specific differentiation program, this program needs to be molecularly accessible. We therefore set out to determine culture conditions for which transient GATA4-mCherry expression led to efficient expression of endogenous GATA6. While in the presence of feeders and 15% serum a 6 h pulse of GATA4-mCherry expression resulted in approx. 10% GATA6-positive cells one day later, this proportion dropped to approx. 1.5% GATA6-positive cells when cells were cultured without feeders in 10% serum (supplementary material Fig. S3A, B), even though GATA4-mCherry was efficiently induced in both conditions (supplementary material Fig. S3C), and cells were positive for the pluripotency marker Nanog (supplementary material Fig. S3A). Next, we pre-cultured cells in 2i + leukemia inhibitory factor (LIF), a condition reported to promote extraembryonic differentiation potential (Morgani et al., 2013), before simultaneous addition of doxycycline and transfer into serum-containing medium. This increased the proportion of GATA6-positive cells induced by a 6 h doxycycline pulse from 11.3 ± 1.8 % (mean ± standard deviation (SD)) for 1 day of pre-culture in 2i + LIF to 51.7 ± 9.8 % for 7 days of pre-culture (Fig. 2A - C). Because the duration of the preculture in 2i + LIF also affected the levels of GATA4-mCherry expression induced by doxycycline (Fig. S4A), we determined the ratio between the fraction of GATA6-positive cells one day after a 6 h doxycycline pulse and the fraction of GATA4-mCherry-positive cells immediately after the pulse as a measure for the efficiency of PrE-like differentiation. This ratio plateaus at approximately 55% after 3 days of preculture in 2i + LIF (Fig. 2C).

**Fig. 2:**
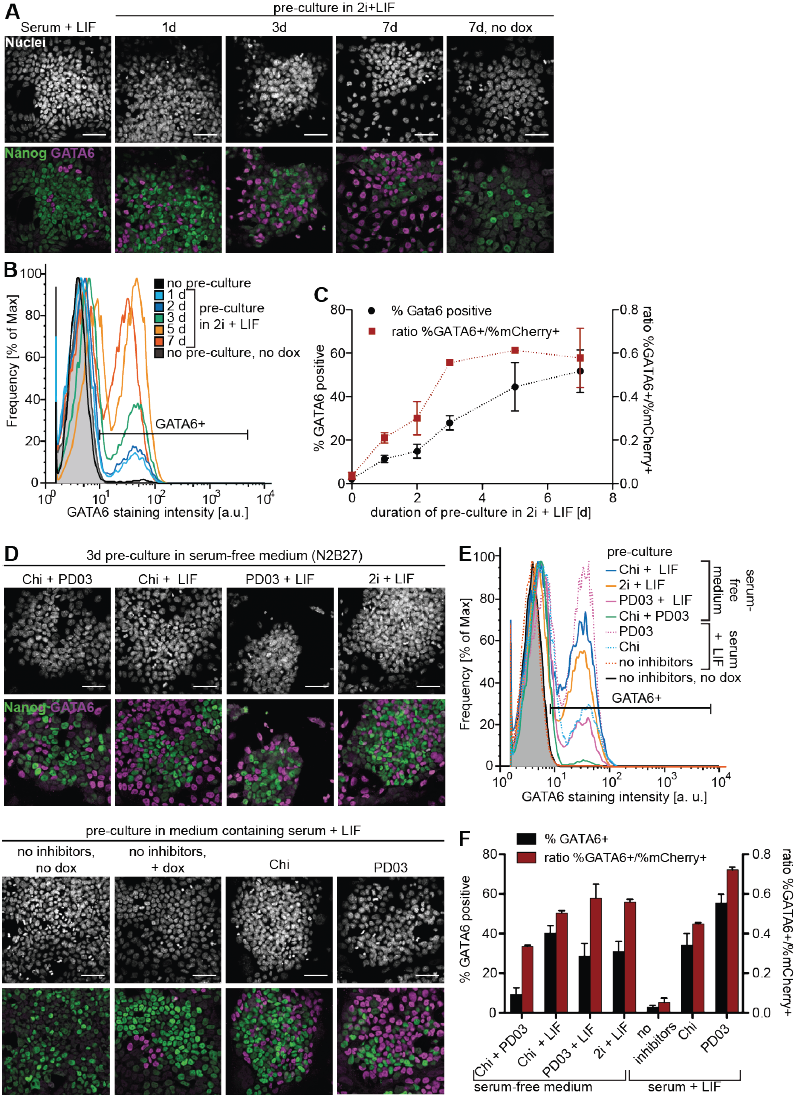
Culture conditions affect responsiveness to GATA4-mCherry expression. ***A*** *Immunostaining of ES cells cultured for indicated times in 2i + LIF before a 6 h pulse of GATA4-mCherry expression followed by a 24 h chase in medium containing serum + LIF. Scale bars, 50 μm.* ***B*** *Flow cytometry of cells treated as in A and stained for GATA6.* ***C*** *Percentage of GATA6-positive cells (black) and ratio of the percentages of GATA6-positive and mCherry-positive cells (red) for different durations of pre-culture in 2i + LIF. Data averaged from 3 (% GATA6-positive) or 2 (ratios) independent experiments, errors bars state standard deviation (SD).* ***D*** *Immunostaining of ES cells cultured for 3 days in the indicated media before a 6 h pulse of GATA4-mCherry expression followed by a 24 h chase in medium containing serum + LIF. Chi, CHIR99021 (3μM). Scale bars, 50 μm.* ***E*** *Flow cytometry of cells treated as in D stained for GATA6.* ***F*** *Average percentage of GATA6-positive cells (black) and ratio of the percentages of GATA6-positive and mCherry-positive cells (red) for different pre-culture media. Data averaged from 3 (% GATA6-positive) or 2 (ratios) independent experiments, errors bars indicate SD.*

To assess the influence of the components of 2i + LIF, we removed each of them from the complete 2i + LIF medium or added them individually to serum-containing medium during 3 days of pre-culture. All conditions led to an increase in the percentage of GATA6-positive cells 24 h after a 6 h doxycycline pulse, albeit to different degrees (Fig. 2D, E, F). The largest proportion of GATA6-positive cells was obtained for pre-culture in serum, LIF and the Mek inhibitor PD0325901 (PD03), (Fig. 2F, supplementary material Fig. S4B). We conclude that inhibition of MAPK signaling prior to the induced expression of GATA factors efficiently restores PrElike differentiation potential in ES cells. For all following experiments, we therefore pre-cultured cells for 3 days in the presence of PD03 in medium containing serum and LIF.

### Transient expression of exogenous GATA4-mCherry induces stable PrE-like differentiation

Having established an experimental regime which induced PrE-like differentiation in ES cells with an efficiency mimicking PrE differentiation in the embryo (Schrode et al., 2014), we next wanted to investigate the stability of the GATA6-positive state and the dynamics with which it evolved. To this end we created a Gata6:H2B-Venus transcriptional reporter in cells carrying the inducible GATA4-mCherry transgene, which faithfully recapitulated GATA6 protein expression between one and three days after the doxycycline pulse (Fig. 3A, B, supplementary material Fig. S5). Transient GATA4-mCherry expression led to a characteristic bimodal distribution of Gata6:H2B-Venus expression (Fig. 3C). Venus expression levels of cells in the Venus^high^ peak were constant between 32 h and 80 h after the end of a 6 h doxycycline pulse (Fig. 3C, D). Furthermore, while cells with intermediate Venus levels progressively disappeared from the distribution (Fig 3C), cells sorted for highest Venus expression levels maintained their fluorescence intensity over several cell divisions for 48h (Fig. 3E). Together, these findings indicate that strong reporter expression marks a stable state in individual cells, and suggests that the decrease in the proportion of Venus^high^ cells is mainly due to reduced proliferation of the Venus^high^ cells compared to the Venus^low^ cells, although we cannot rule out that undifferentiated cells induce reversion of Venus^positive^ cells in unsorted populations.

**Fig. 3:**
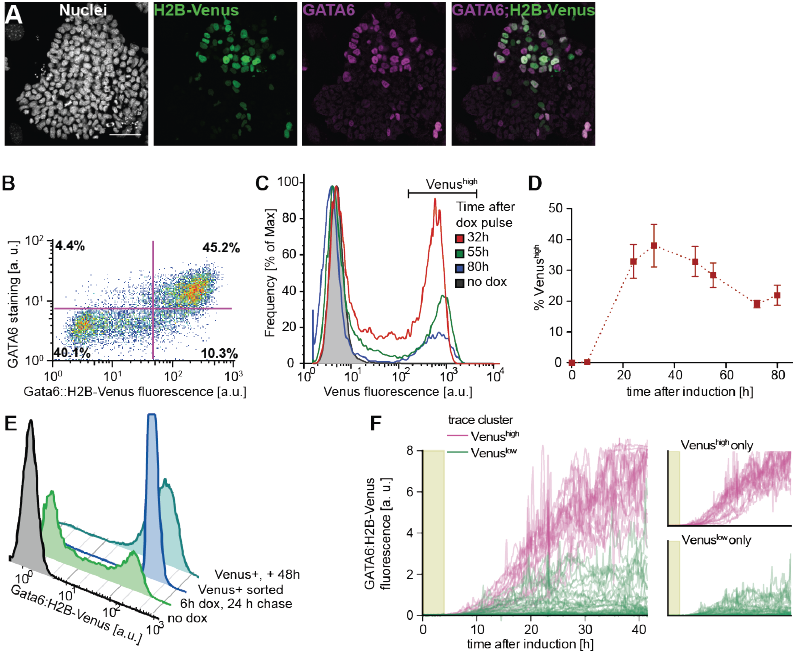
Transient GATA4-mCherry expression induces stable expression of PrE marker genes. ***A*** *Immunostaining for Venus and GATA6 protein in Gata6:H2B-Venus reporter cells 24 h after a 6 h pulse of doxycycline-induced GATA4-mCherry expression. Scale bar, 50 μm.* ***B*** *Flow cytometry of Gata6-reporter cells treated as in* ***A***. ***C*** *Flow cytometry detecting Gata6:H2B-Venus expression at indicated time-points after a 6 h pulse of GATA4-mCherry induction.* ***D*** *Percentages of Venus^high^ cells at different times after a 6 h doxycycline pulse. Data points represent mean* ± *SD from three independent experiments.* ***E*** *Flow cytometry of Venus^high^ cells sorted 18 h after the end of a 6 h doxycycline pulse (dark blue), cultured for 48 h and analyzed for Venus expression.* ***F*** *H2B-Venus fluorescence intensity values of individual Gata6-reporter cells tracked in time-lapse movies during and following a 6 h doxycycline pulse (shaded area). Color-code is informed by hierarchical clustering based on H2B-Venus expression levels. Small panels on the right show traces for each cluster separately. See also supplementary material Fig. S6 and Movie 1.*

Finally, to follow the dynamics of Gata6:H2B-Venus expression over time in individual cells, we performed time-lapse microscopy and tracking of reporter cells (supplementary material Movie 1). Clustering of traces according to H2B-Venus expression levels identified two distinct classes of cells (Fig. 3F, supplementary material Fig. S6). Expression traces corresponding to these two classes were separated throughout the experiment. While some cells with intermediate Venus levels reverted to a Venus-negative state, consistent with the depletion of this population that we had observed by flow cytometry, cells with highest Venus expression remained in this class throughout the experiment (Fig. 3F). These results show that a transient GATA4-mCherry input elicits stable expression of one of two mutually exclusive expression programs, suggesting the system behaves as an irreversible switch with two stable states.

### A threshold level of GATA4-mCherry controls PrE-like differentiation

We then asked whether the flipping of the bistable switch that we had identified depended on the expression levels of the doxycycline-induced GATA4-mCherry protein. Varying the duration and levels of GATA4-mCherry exposure by applying doxycycline pulses of different length (supplementary material Fig. S7) smoothly tuned the proportion of Gata6:H2B-Venus^high^ cells (Fig. 4A, B). Furthermore, we observed more differentiating GATA6-positive cells and fewer undifferentiated Nanog-positive cells in populations that had been sorted for high GATA4-mCherry expression levels immediately after the doxycycline pulse compared to populations sorted for low GATA4-mCherry expression (Fig. 4C, D, supplementary material Fig. S8). Together, this suggests that GATA4-mCherry expression levels control the proportion of cells undergoing PrE-like differentiation.

**Fig. 4:**
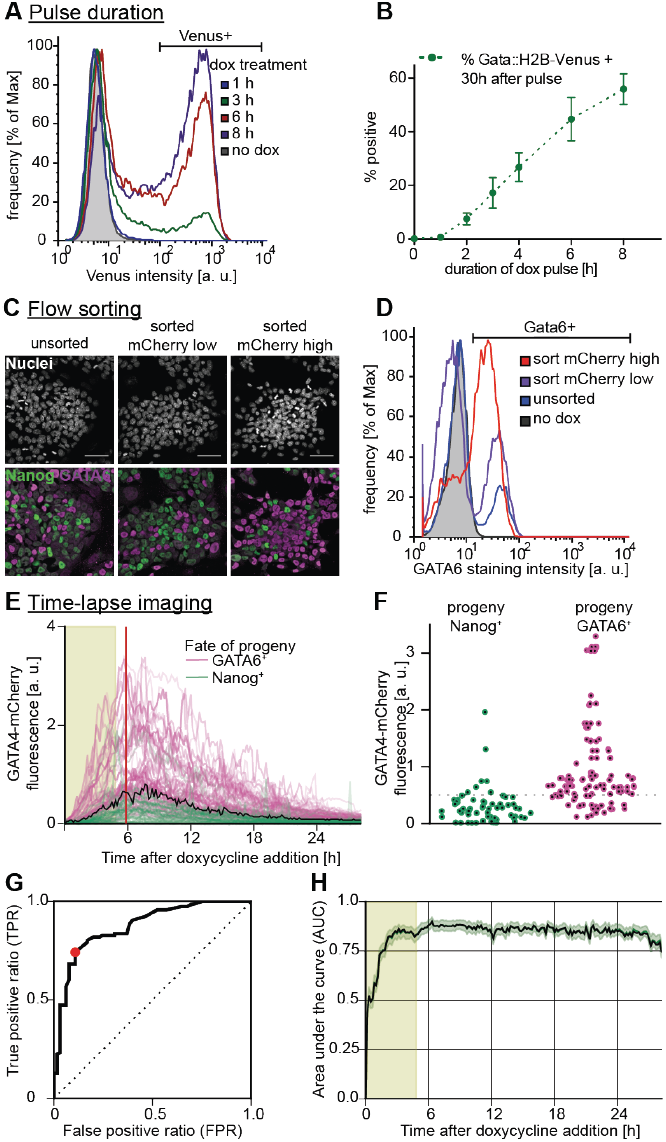
A GATA4-mCherry threshold dose determines PrE-like differentiation. ***A*** *Flow cytometry for Gata6:H2B-Venus fluorescence one day after transient GATA4-mCherry expression triggered by doxycycline pulses of indicated lengths.* ***B*** *Quantitative analysis of results from* ***A****. Data points show mean and SD from three independent experiments.* ***C*** *Cells sorted for low (middle) and high (right) GATA4-mCherry expression levels after a 6 h doxycycline pulse and immunostained 30 h after re-plating. Unsorted control is on the left. Scale bar, 50 μm.* ***D*** *Flow cytometry of cells treated as in* ***C*** *stained for GATA6 expression. Purification increases the proportion of differentiating cells in both sorted pools compared to the unsorted control, and a larger proportion of GATA4-mCherry^high^ cells activate GATA6 expression compared to GATA4-mCherry^low^ cells.* ***E*** *GATA4-mCherry fluorescence traces of cells filmed during and after a 6 h doxycycline pulse. Color-code of individual traces is informed by immunostaining for Nanog and GATA6 at the end of the time-lapse. Area shaded in green indicates duration of doxycycline pulse, red bar indicates time point analyzed in* ***F***, ***G****, and black curve indicates optimal threshold calculated by ROC. See also supplementary material Movie 2 and Fig. S9.* ***F*** *GATA4-mCherry fluorescence intensity of cells from the experiment shown in E at a single time-point (red vertical line in* ***E****). Optimal threshold to predict fate choice estimated by ROC analysis is indicated by a black line.* ***G*** *ROC curve for the time-point shown in* ***F****. The optimal threshold maximizing the difference between the true positive and false positive prediction rate (TPR and FPR) is indicated by a red dot.* ***H*** *Area under the curve (AUC) values from ROC analysis in all time frames of the experiment shown in* ***E****. Error margin indicates standard deviation determined by bootstrapping (n = 1000).*

To correlate GATA4-mCherry input levels more precisely with subsequent fate choice in single cells, we performed time-lapse imaging of GATA4-mCherry-inducible cells during and after a doxycycline pulse, followed by immunostaining for Nanog and GATA6 (supplementary material Movie 2, Fig. S9A, B). We found that most cells with GATA6-positive progeny had experienced higher GATA4-mCherry expression levels than cells with Nanog-positive progeny (purple and green datapoints in Fig. 4E, F, and in supplementary material Fig. S9C). We used receiver operating characteristics (ROC) analysis (Fawcett, 2006; for details see supplementary material) to assess how well the two classes of cells could be separated by a threshold value of GATA4-mCherry expression. Plotting the ratio of correctly and incorrectly separated events over the total number of cells (true positive ratio TPR, and false positive ratio, FPR) for varying threshold values gives a characteristic curve for a single time-point (Fig. 4G); the larger the area under this curve (AUC), the better the differentiation outcomes can be separated or predicted from GATA4-mCherry expression levels. The AUC increased quickly upon doxycycline addition and reached a plateau between 0.8 and 0.9 after approx. 3 h (Fig. 4H). Similar results were obtained when we used cumulative instead of instantaneous GATA4-mCherry expression measurements (not shown), suggesting that non-systematic measurement errors are not a major limit of predictive power. The optimal prediction threshold that maximizes the difference between TPR and FPR tracked the expression dynamics of the GATA4-mCherry protein throughout the experiment (black line in Fig. 4E, supplementary material Fig. S10A, B). Using this threshold, more than 80% of all fate decisions could be correctly predicted based on the GATA4-mCherry classifier (supplementary material Fig. S10C). This predictability of fate choice by GATA4-mCherry expression levels suggests this transcription factor is a dominant input into the decision in ES cells.

### FGF/MAPK signaling modulates the proportion of cells with PrE-like differentiation

In the mouse embryo, both GATA factors and FGF/MAPK signaling are required to establish PrE identity. Having shown above that inhibiting MAPK signaling prior to doxycycline-induced GATA expression increases the proportion of cells with PrE-like differentiation, we next wanted to test how MAPK signaling affected the decision to embark on PrE-like differentiation during and after the GATA pulse. MAPK activity required for PrE-like differentiation was almost completely saturated in serum-free medium, possibly through autocrine FGF signaling (supplementary material Figs S11, S12), prompting us to tune the levels of Erk phosphorylation following removal of the pre-culture medium with subsaturating doses of PD03 (Fig. 5A). Partial inhibition of MAPK signaling during the 6 h doxycycline pulse and the 24 h chase period reduced the fraction of GATA6 positive cells, but not the expression levels of GATA6 in individual differentiated cells (Fig. 5B, C), with a quasi-linear relationship between the level of Erk phosphorylation and the percentage of differentiating cells (Fig. 5D). We obtained similar results using the FGF receptor inhibitor PD173074 (supplementary material Fig. S12), indicating that most of the MAPK activity relevant for PrE-like differentiation of ES cells is triggered by FGF ligands, consistent with literature reports (Kunath et al., 2007). FGF/MAPK signaling levels therefore control the fraction of cells that embark on the PrE-like differentiation path.

**Fig. 5:**
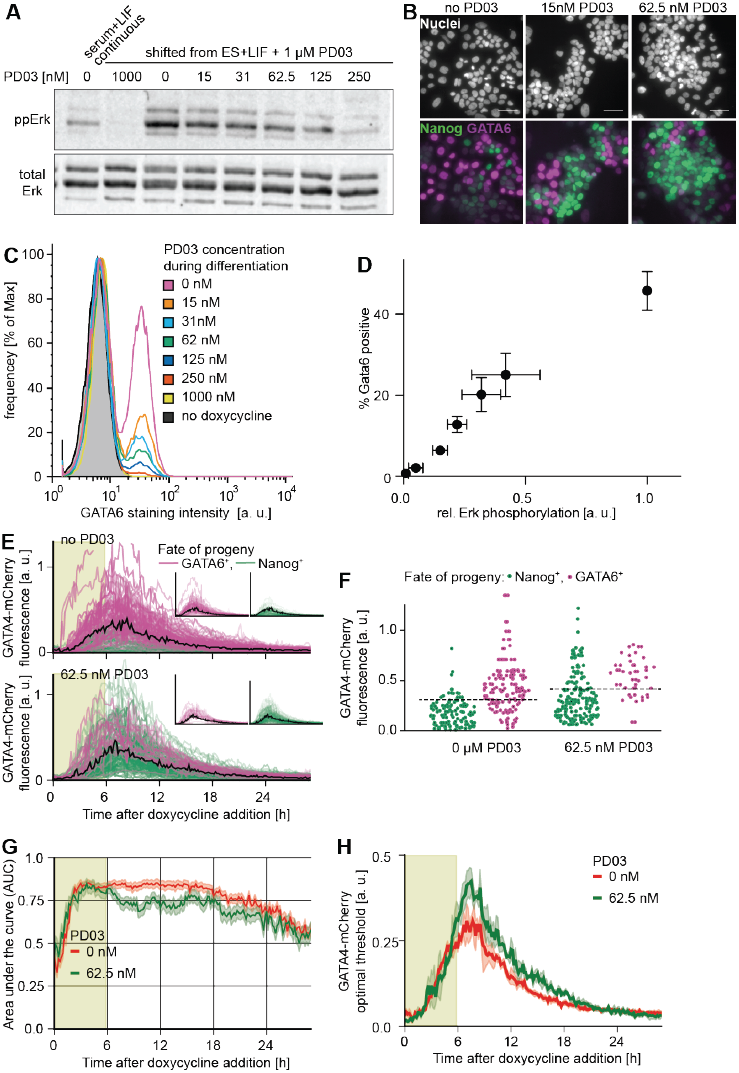
MAPK signaling controls the proportion of cells with PrE-like differentiation. ***A*** *Immunoblot detecting phosphorylated Erk (top) and total Erk (bottom) in cells grown for 3 d in the presence of serum and 1 μM PD03 1 h after transfer into medium containing indicated amounts of PD03.* ***B*** *Immunostaing of ES cells after a 6 h pulse of doxycycline-induced GATA4-mCherry expression and 24 h of differentiation in the indicated concentrations of PD03. Scale bar, 50 µm.* ***C*** *Flow cytometry of GATA6 expression in cells treated as in* ***B***. ***D*** *Plot of relative Erk phosphorylation levels versus percentage of GATA6-positive cells for different concentrations of PD03. Data points show mean and SD from three independent experiments per condition.* ***E*** *GATA4-mCherry fluorescence traces of cells filmed during and after a 6 h doxycycline pulse in the absence of PD03 (top) or with 62.5 nM PD03 (bottom). Color-code of individual traces is informed by immunostaining for Nanog and GATA6 at the end of the time-lapse. Shaded area indicates presence of doxycycline, black trace indicates optimal threshold estimated by ROC, and insets show traces for cells with GATA6- and Nanog-positive progeny separately.* ***F*** *GATA4-mCherry fluorescence intensity of cells from the experiment shown in* ***E*** *at a single time-point 1 h after removal of doxycycline in the absence of PD03 (left) and in 62.5 nM PD03 (right). G AUC values determined by ROC analysis of the dataset shown in* ***E*** *for no PD03 (red) and 62.5 nM PD03 (green). AUC values decay after approx. 20 h due to the very low levels of GATA4-mCherry expression in this experiment. Error margin indicates standard deviation determined by bootstrapping (n=1000).* ***H*** *The optimal threshold to predict differentiation from GATA4-mCherry expression in the absence of PD03 (red) and in the presence of 62.5 nM PD03 (green) increases with decreasing MAPK signaling. Error margins indicate standard deviation determined by bootstrapping (n=1000).*

To investigate how partial Mek inhibition affected the GATA4-mCherry threshold required for PrE-like differentiation, we performed time-lapse imaging and cell tracking for maximal and reduced MAPK signaling in parallel (Fig. 5E, F), using a PD03 concentration that led to a significant reduction of the number of differentiating cells without inducing cell death (supplementary material Movie 3). ROC analysis gave similar AUC values for both conditions, indicating that differentiation can be predicted based on GATA4-mCherry expression levels with similar confidence at different signaling levels (Fig. 5G). However, the optimal prediction threshold was consistently increased upon partial Mek inhibition (Fig. 5F, H). We conclude that MAPK signaling levels set the GATA4-mCherry threshold dose required to trigger differentiation.

We noticed that the distribution of GATA4-mCherry expression levels in differentiating and non-differentiating cells changed upon partial inhibition of signaling (Fig. 5E, F). In addition to setting the transcription factor threshold dose, partial Mek inhibition therefore appears to modulate heterogeneities in the population that affect PrE-like differentiation.

### A simple mutual repression circuit recapitulates the experimentally observed gene expression dynamics

To gain insights into the formal nature of the interactions between signaling and transcriptional regulators we then sought to identify the minimal circuit model of the components of the decision machinery that would recapitulate our data. The irreversible, switch-like behavior of our system indicates the presence of positive feedback in the underlying genetic network. Because Nanog directly represses Gata6 (Singh et al., 2007), and GATA expression led to rapid repression of Nanog in our system, we chose a network of two mutually repressive nodes, Gata and Nanog, as a minimal system with net positive feedback to formalize a bistable genetic switch (Cherry and Adler, 2000; Plahte et al., 1995; Snoussi, 1998; Thomas, 1981) (Fig. 6A, see supplementary material for a detailed description of the model). This system is described by two coupled ordinary differential equations that account for the dynamics of Nanog (N) and endogenous GATA (G) as markers for the Epi and PrE programs in individual cells, respectively:

**Fig. 6:**
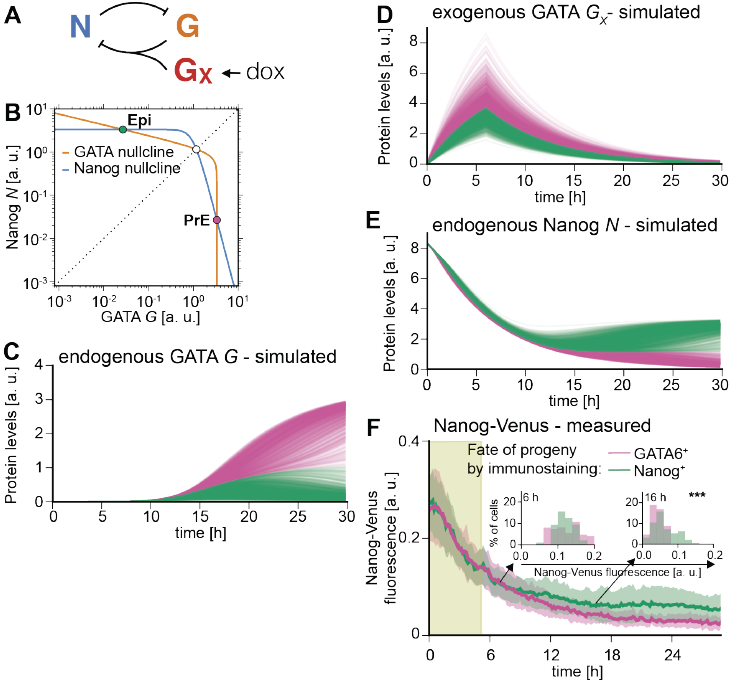
A simple mutual repression circuit recapitulates experimentally observed expression dynamics. ***A*** *Connectivity of the mutual repression circuit used in dynamic simulations.* ***B*** *Phase portrait depicting the autonomous dynamics of the system in* ***A****. Intersections of the Nanog nullcline (blue) and the GATA nullcline (orange) define three steady states, two of which are stable and correspond to the Nanog^high^, GATA^low^ Epi state (green dot) or the Nanog^low^, GATA^high^ PrE state (purple).* ***C***, ***D***, *E Simulated time traces of endogenous GATA G (****C****), exogenous GATA4-mCherry G_x_ (****D****), and Nanog expression (****E****). Color code is informed by final GATA expression levels in* ***C*** *(GATA^htgh^: purple, and GATA^low^: green), and corresponding traces have same color in* ***C***, ***D*** *and* ***E***. ***F*** *Average Nanog-Venus expression levels in cells carrying a Nanog-Venus translational reporter during and after 6 h of doxycycline-induced GATA4-mCherry expression. Cells were classified as GATA6-positive and Nanog-positive by immunostaining after the time-lapse. Presence of doxycycline is indicated by shaded green rectangle. Lines and shaded areas indicate mean Nanog-Venus fluorescence levels and standard deviation. Insets show histograms for distributions of Nanog-Venus fluorescence in the two classes of cells at 6 h and 16 h after the start of recording;* *** *indicates p ≤ 0.0001 (Mann-Whitney’s U-test). See also supplementary material Movie 4.*

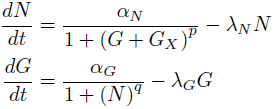

A third equation modxls the externally supplied pulse of GATA (*G_x_*) that drives the endogenous circuit:

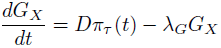

To reflect the experimentally observed heterogeneous expression of exogenous GATA factors, we varied the maximum transcription rate *D* of exogenous GATA between cells (for details see supplementary material). This was the only source of cell-to-cell variability in our model. As initial conditions, we endowed cells with high levels of Nanog and no GATA to reflect pre-culturing in the presence of PD03.

To assess the dynamics of the endogenous circuit described by this model we plotted the nullclines *dG/dt=0* and *dN/dt=0* for the specific set of parameters used. Two of the three equilibrium states defined by the intersections of the nullclines are stable and correspond to the fully differentiated GATA-positive and the undifferentiated Nanog-positive state, respectively (Fig. 6B). A boundary in the phase space (dashed line in Fig. 6B) separates the combinations of GATA and Nanog levels which will evolve into the fully differentiated GATA-positive state from those that lead to the undifferentiated Nanog-positive state. To induce PrE-like differentiation, the exogenous GATA input has to exceed the threshold required to move cells across this boundary by sufficiently repressing Nanog and allowing for endogenous GATA expression. For the chosen parameter set, simulated time traces of individual cells closely resembled the experimentally observed expression dynamics of the endogenous Gata6 gene (Fig. 6C, compare to Fig. 3G), and exogenous GATA4-mCherry (Fig. 6D, compare to Fig. 4E), suggesting this simple mutual repression circuit is sufficient to capture essential dynamics of the experimental system.

To further test the model, we compared the dynamics of Nanog expression *in silico* and *in vivo.* In model simulations, Nanog expression levels first decreased rapidly in all cells from the initial conditions chosen to represent the effects of the pre-culture regime towards lower steady state levels, before differences in Nanog expression levels in differentiating and non-differentiating cells became apparent (Fig. 6E). These expression dynamics were observed experimentally in cells expressing a translational Nanog fusion reporter (Filipczyk et al., 2013) following transient expression of GATA4-mCherry (supplementary material Movie 4, Fig. 6F). This agreement between Nanog expression dynamics *in silico* and *in vivo* further supports the idea that a simple mutual repression circuit is sufficient to capture the dynamics of the system.

### Inhibition of the Epi program by MAPK signaling controls the proportion of cells with PrE-like differentiation

To pinpoint the main mechanism by which FGF/MAPK signaling controls the fraction of cells with PrE-like differentiation, we considered two simple extensions of the model, one in which FGF/MAPK signaling promotes expression of the PrE program (Fig. 7A), and an alternative model in which signaling inhibits the Epi program (Fig. 7B). In both cases a reduction of signaling led to a simulated increase in the number of cells in the Nanog-positive peak and a decrease of cells in the GATA-positive peak (Fig. 7A, B, middle). However, expression levels in the respective positive peaks changed in distinct ways depending on the type of signaling input (Fig. 7A, B, middle). Estimating changes in GATA6 and Nanog expression levels upon partial Mek inhibition from flow cytometry data showed that GATA6 expression levels in individual cells remained approximately constant in the presence of different doses of PD03, while the Nanog-Venus positive peak consistently shifted to higher expression levels with lowered signaling (Fig. 7C, D, supplementary material Fig. S13). While not ruling out a more complex integration of FGF/MAPK signaling into the regulatory circuit underlying PrE-like differentiation, these results suggest that a major route by which FGF/MAPK signaling controls the fraction of cells with PrE-like differentiation is through inhibition of the Epi-specific gene expression program. This conclusion is further supported by a recent report showing direct inhibition of Nanog expression by FGF/MAPK signaling mediated by chromatin modifications (Hamilton and Brickman, 2014).

**Fig. 7:**
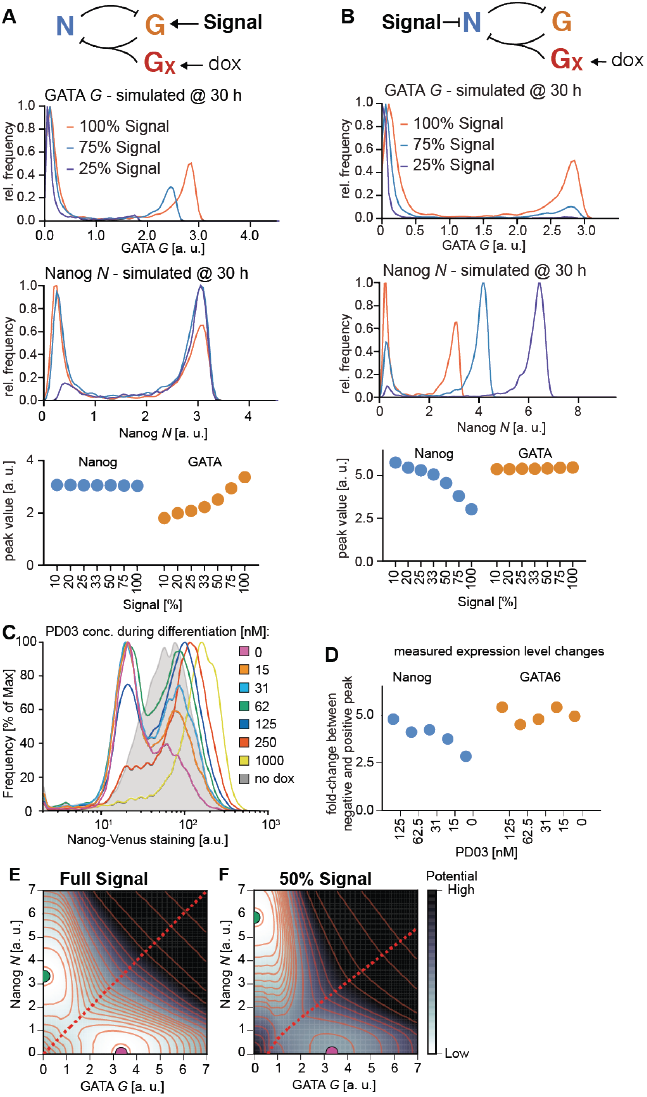
MAPK signaling controls the proportion of PrE-like cells through inhibition of the Epi program. ***A, B*** *Top: Modes of signaling interactions tested in the model.* ***A****: signaling promotes GATA production,* ***B****: Signaling inhibits Nanog production. Middle: Simulated histograms and location of histogram peaks for GATA and Nanog expression at varying signaling levels (t = 30 h; n = 2000).* ***C*** *Flow cytometry of Nanog-Venus reporter cells stained with antibodies directed against the Venus tag 30 h after a 6 h doxycycline pulse. This method detected a bimodal distribution of Nanog-Venus-positive and –negative cells with a higher signal-to-noise ratio than direct staining for Nanog. Traces are smoothened to show positions of peaks more clearly.* ***D*** *Fold-change of the Nanog-Venus-positive (blue) and GATA6-positive (orange) peaks relative to the respective negative peaks estimated from the flow cytometry experiments in* ***C*** *(Nanog-Venus) and Fig. 5* ***C*** *(GATA6) for different concentrations of PD03. See supplementary material Fig. S13 for details on estimation of peak positions.* ***E***, ***F*** *Quasi-potentials derived from the autonomous dynamics of the system in* ***B*** *for two signaling levels. Darker regions have higher potential, i.e. faster dynamic changes. The dashed red line indicates the separatrix (ridge) separating the basins of attraction of the Nanog^high^, GATA^low^ Epi fate (green dot) and the Nanog^low^, GATA^high^ PrE fate (purple dot). Lowered signaling (****F****) shifts and bends the separatrix, thereby changing the relative sizes of the two basins of attraction.*

Finally, we sought to develop a visual representation of the system’s dynamics in different signaling regimes by estimating the path-integral quasi-potential surfaces (Bhattacharya et al., 2011) of the system for two different signaling levels (Fig. 7E, F). This representation highlights two basins of attraction corresponding to the Nanog-positive state and the GATA positive state, respectively. A reduction in signaling bends the ridge that separates the basins towards the GATA-positive state, making its basin of attraction narrower and shallower relative to that of the Nanog-positive state (Fig. 7E, F).

We conclude that a simple mutual repression circuit is sufficient to capture the dynamic hallmarks of the Epi-versus-PrE fate decision, and that through repression of the Epi program, FGF/MAPK signaling sets the relative sizes of the basins of attraction corresponding to the two fates defined by this circuit, allowing signaling to regulate the proportions of cells adopting either fate.

## Discussion

Here we have used engineered mouse ES cell lines to study the mechanism underlying the decision between the Epi and the PrE fate. Our experimental system allowed us to modulate and measure quantitatively the transcription factor and signaling inputs into the decision, and to follow the dynamics of the decision at the level of single cells. We have uncovered two successive functions of MAPK signaling in the ES cell system: Before the induced expression of GATA factors, inhibition of MAPK signaling is required to make the PrE-like differentiation program accessible in ES cells. Once exogenous GATA factors are expressed, MAPK signaling is required to execute the decision of PrE-like differentiation. The Epi-versus-PrE differentiation event displays hallmarks of an irreversible bistable switch, as coexpression of determinants of the Epi and the PrE fate resolves into one of two mutually exclusive stable states characterized by Nanog and GATA expression, respectively. We detect a well-defined threshold level of exogenous GATA factor expression required to flip this switch and induce differentiation, and find that MAPK signaling sets this threshold dose. This decision is therefore a strongly regulated process that is largely determined by few well-defined transcriptional and signaling inputs.

### The accessibility of the PrE program depends on ES cell culture conditions

We find that cells cultured in the presence of serum are refractory to PrE-like differentiation upon induced GATA expression, but responsiveness to doxycycline-induced GATA factors can be restored by extended exposure to GSK3 or Mek inhibitors, e.g. in 2i medium. One interpretation of this finding is that ES cells cultured in serum are strongly biased towards embryonic fates, and as a consequence have blocked the PrE-like differentiation program. In line with this idea, ES cells grown in the presence of serum display higher levels of repressive chromatin marks on a subset of promoters, including the Gata6 promoter, than cells grown in 2i + LIF (Marks et al., 2012). Furthermore, the transcriptome of ICM cells resembles more closely that of ES cells cultured in 2i medium than that of ES cells cultured in serum (Boroviak et al., 2014). This suggests that pre-culture in 2i brings ES cells to a molecular state mirroring that of early ICM cells, from which, upon induced GATA expression, the decision between the Epi and the PrE fate can be taken similar to the situation in the embryo.

### Extraembryonic fate choice is determined by the output of a simple mutual repression circuit

Our finding that precise measurements of GATA4-mCherry expression levels allow prediction of fate decisions in individual ES cells with high confidence before endogenous fate markers appear led us to formulate a minimal genetic circuit model with deterministic regulation to formalize the mechanism of the decision process. Our model solely consists of mutually repressive interactions between the Epi- and the PrE-like program, modulated by a repressive input of FGF/MAPK on the Epi program (Fig. 7B). This is sufficient to recapitulate the experimentally observed dynamics of lineage marker expression, to model bistable behavior, and to formalize our finding that the role of MAPK signaling is to set a GATA threshold required for PrE-like differentiation. Our minimal model is a subnetwork of a more complex model for the Epi-versus-PrE fate decision recently developed by Bessonard et al. (2014). Bessonard’s model posits an additional positive input of FGF/MAPK signaling onto Gata expression, and contains positive autoregulatory feedback loops centered on both Nanog and Gata, which endow the dynamic system with a third stable state of Nanog and GATA co-expression. This allowed Bessonard et al. to simulate both the establishment and the resolution of the co-expression state in a single model. Focusing on the resolution of the co-expression state, our data suggest that the additional links of Bessonard’s model are not required to explain the dynamics of this phase of the decision. It remains however possible that positive autoregulation of the Epi and PrE programs and a positive input of FGF/MAPK signaling on Gata expression fine-tune the response of cells during this stage of the decision process.

We note that not all cells abide to the GATA4-mCherry threshold that best predicts PrE-like differentiation. This may reflect persistent heterogeneous chromatin configurations that block PrE-like differentiation in individual cells, or be a consequence of heterogeneous MAPK signaling amongst ES cells. Signaling heterogeneities have been detected in other cell lines (Albeck et al., 2013; Aoki et al., 2013), and we expect they will be functionally relevant for PrE-like differentiation of ES cells.

### Integration of signaling into the mutual repression circuit serves to balance proportions of cell fates in developing tissues

The mathematical model of a mutual repression circuit has previously been applied to describe the dynamics of the switch between the lysogenic and lytic phases of the lifecycle of bacteriophage lambda (Ptashne, 2004), and a genetically engineered toggle switch circuit in E. coli (Gardner et al., 2000). Our work is one of the first experimentally supported examples demonstrating that this network can be used to formalize the decision between two fates during mammalian development. Extending the model with a signaling input allows for dynamic control of the sizes of the basins of attraction corresponding to the different states of the bistable system. The mammalian preimplantation embryo may harness this property to balance the proportion of Epi and PrE cells. The initial expression of transcriptional regulators driving lineage choice is stochastic, possibly due to the mechanisms that control gene expression in the early embryo (Dietrich and Hiiragi, 2007; Ohnishi et al., 2014), and the resulting heterogeneous distributions of transcription factor concentrations in ICM cells bias cells towards specific fates (Xenopoulos et al., 2015). It has been shown that lineage commitment occurs non-synchronously in the cells of the ICM, and that the first cells to commit are fated towards the epiblast (Grabarek et al., 2012). Because Epi cells produce Fgf4 (Frankenberg et al., 2011; Nichols et al., 1998), Fgf4 levels will reflect the number of Epi-committed cells and act on the as-yet uncommitted cells. By modulating the bistable switch operating in these cells, this process may ultimately place the appropriate number of uncommitted cells in the basin of attraction corresponding to the PrE fate. FGF/MAPK signaling might thus acts as feedback mechanism to balance proportions of two distinct cell fates in populations (Lander et al., 2009). It will be interesting to see whether this new principle applies to differentiation decisions beyond those in the preimplantation embryo.

## Experimental Procedures

### ES cell culture and genetic manipulation

For genetic engineering, ES cells were grown on mitotically inactivated mouse embryonic fibroblasts in Knockout DMEM (Gibco) supplemented with 15% fetal bovine serum (FBS), 50 µM β-mercaptoethanol, glutamax, non-essential amino acids and 1 µg/ml LIF. After line derivation, feeders were removed by serial passaging, and cells were maintained on gelatin coated dishes in GMEM-based medium supplemented with 10% FBS, sodium pyruvate, 50 µM β-mercaptoethanol, glutamax, non-essential amino acids and LIF.

Serum-free media were based on N2B27 (NDiff 227, Stem Cells) and supplemented with 3 μM CHIR99021, 1μM PD0325901 and 1 μ/ml LIF to give 2i + LIF. For the experiments described in supplemenatary material Fig. S11, N2B27 was supplemented with 10 ng/ml BMP4 (R&D), 1 μ/ml LIF and 1 μ/ml heparin.

All cell lines used in this study were based on the KH2 ES cell line (Beard et al., 2006). Engineering of ES cells is described in more detail in the supplementary material. Transgene expression was induced by adding 500 ng/ml doxycycline to the culture medium.

### Immunocytochemistry

Cells for immunocytochemistry were grown on ibidi μ-slides and stained as described in Kalmar et al., 2009. Primary antibodies were anti-Nanog (eBiosciences, 1:200), anti-FLAG (Sigma M2, 1:1000), anti-GATA6 (R&D, 1:200) and anti-GATA4 (Santa Cruz, 1:200). Detection was performed using Alexa-Fluor conjugated secondary antibodies (Molecular Probes). Nuclei were visualized using Hoechst 33342 dye (Molecular Probes). Cells were images on a Zeiss LSM700 confocal microscope with a 40x oil immersion lens (NA 1.2).

### Flow cytometry

Cells for flow cytometry were trypsinized and either analyzed immediately or fixed for 15 minutes in 3% PFA/PBS. Intracellular antigens were stained in suspension using the same primary and secondary antibodies as used for immunostaining. mCherry fluorescence was measured on a BD Fortessa Flow cytometer, all other flow cytometric analysis was performed using a Beckman Coulter CyAn ADP analyzer. Cell sorting was done on a Beckman Coulter MoFlo. To estimate peak positions, histograms were smoothened, followed by detection of local maxima with custom-written Python scripts.

### Immunoblotting

For immunoblotting, cells were lysed in RIPA buffer and lysates were separated on polyacrylamide gels before transfer to nitrocellulose membranes. Antibodies used were anti-ppErk (Sigma M9692) and anti-Erk1/2 (Millipore 06-182) at 1:500 dilution. Detection was performed using fluorescently labeled secondary antibodies and scanning in a LI-COR Odyssey system. Intensities of bands were quantified in ImageStudio (LI-COR).

### Time-lapse imaging and cell tracking

Time-lapse imaging was performed in DMEM based medium without phenol red, supplemented as detailed above. We used a Zeiss Axiovert M200 microscope equipped with a SOLA LED light source, an Andor iXON Ultra 888 EMCCD camera and a heated stage with CO_2_ supply. Hardware was controlled by MicroManager software (Edelstein et al., 2001). Time-lapse movies were acquired using a 40x long-working distance lens. See Supplemental Experimental Procedures for details on image analysis.

### Mathematical modeling

Numerical simulations of the model were implemented in Python language. Parameter values used in the simulations are given in supplementary material Table S1. For details on the model see supplementary material.

## Author contributions

CS, PR and AMA conceived the study; CS performed experiments; CS, PR and JPM analyzed the data; PR developed the mathematical model; CS wrote the manuscript with input from PR and AMA; all authors approved the manuscript.

## Acknowledgements

We thank K. Niakan for providing cell lines, and K. Anastassiadis, B. Rosen, C. Lindon and A.-K. Hadjantonakis for sharing reporter constructs. L. Filipkova, N. S. Ly and R. Broome provided technical assistance, and L. Garcia-Perez, C. Vintiner and J. Padua helped with cell tracking. We are grateful to J. Nichols and A.-K. Hadjantonakis for insightful discussions, and C. Pina, J. de Navascués, S. Munoz-Descalzo, C. Brimson, J. Garci-Ojalvo and A. Oates for helpful comments on earlier versions of this manuscript. Work in the Martinez Arias lab was funded by an ERC investigator grant. CS was the recipient of an EMBO long-term fellowship, and CS and PR were supported by a Marie Curie fellowship.

## References

Albeck, J. G., Mills, G. B. and Brugge, J. S. (2013). Frequency-modulated pulses of ERK activity transmit quantitative proliferation signals. Molecular Cell 49, 249–261.

Aoki, K., Kumagai, Y., Sakurai, A., Komatsu, N., Fujita, Y., Shionyu, C. and Matsuda, M. (2013). Stochastic ERK activation induced by noise and cell-to-cell propagation regulates cell density-dependent proliferation. Molecular Cell 52, 529–540.

Beard, C., Hochedlinger, K., Plath, K., Wutz, A. and Jaenisch, R. (2006). Efficient method to generate single-copy transgenic mice by site-specific integration in embryonic stem cells. Genesis 44, 23–28.

Beddington, R. S., and Robertson, E. J. (1989). An assessment of the developmental potential of embryonic stem cells in the midgestation mouse embryo. Development 105, 733–737.

Bessonnard, S., De Mot, L., Gonze, D., Barriol, M., Dennis, C., Goldbeter, A., Dupont, G. and Chazaud, C. (2014). Gata6, Nanog and Erk signaling control cell fate in the inner cell mass through a tristable regulatory network. Development 141, 3637–3648.

Bhattacharya, S., Zhang, Q. and Andersen, M. E. (2011). A deterministic map of Waddington’s epigenetic landscape for cell fate specification. BMC Syst Biol 5, 85.

Boroviak, T., Loos, R., Bertone, P., Smith, A. and Nichols, J. (2014). The ability of inner-cell-mass cells to self-renew as embryonic stem cells is acquired following epiblast specification. Nature Cell Biology 16, 516–528.

Chambers, I., Silva, J., Colby, D., Nichols, J., Nijmeijer, B., Robertson, M., Vrana, J., Jones, K., Grotewold, L. and Smith, A. (2007). Nanog safeguards pluripotency and mediates germline development. Nature 450, 1230–1234.

Cherry, J. L., and Adler, F. R. (2000). How to make a biological switch. J Theor Biol 203, 117–133.

Cho, L. T. Y., Wamaitha, S. E., Tsai, I. J., Artus, J., Sherwood, R. I., Pedersen, R. A., Hadjantonakis, A.-K. and Niakan, K. K. (2012). Conversion from mouse embryonic to extra-embryonic endoderm stem cells reveals distinct differentiation capacities of pluripotent stem cell states. Development 139, 2866–2877.

Dietrich, J.-E., and Hiiragi, T. (2007). Stochastic patterning in the mouse preimplantation embryo. Development 134, 4219–4231.

Edelstein, A., Amodaj, N., Hoover, K., Vale, R. and Stuurman, N. (2001). Computer Control of Microscopes Using μManager. Hoboken, NJ, USA: John Wiley & Sons, Inc.

Fawcett, T. (2006). An introduction to ROC analysis. Pattern Recognition Letters 27, 861–874.

Filipczyk, A., Gkatzis, K., Fu, J., Hoppe, P. S., Lickert, H., Anastassiadis, K. and Schroeder, T. (2013). Biallelic expression of nanog protein in mouse embryonic stem cells. Cell Stem Cell 13, 12–13.

Frankenberg, S., Gerbe, F., Bessonnard, S., Belville, C., Pouchin, P., Bardot, O. and Chazaud, C. (2011). Primitive endoderm differentiates via a three-step mechanism involving Nanog and RTK signaling. Dev Cell 21, 1005–1013.

Fujikura, J., Yamato, E., Yonemura, S., Hosoda, K., Masui, S., Nakao, K., Miyazaki Ji, J.-I. and Niwa, H. (2002). Differentiation of embryonic stem cells is induced by GATA factors. Genes Dev 16, 784–789.

Gardner, T. S., Cantor, C. R. and Collins, J. J. (2000). Construction of a genetic toggle switch in Escherichia coli. Nature 403, 339–342.

Grabarek, J. B., Zyzynska, K., Saiz, N., Piliszek, A., Frankenberg, S., Nichols, J., Hadjantonakis, A.-K. and Plusa, B. (2012). Differential plasticity of epiblast and primitive endoderm precursors within the ICM of the early mouse embryo. Development 139, 129–139.

Hamilton, W. B., and Brickman, J. M. (2014). Erk Signaling Suppresses Embryonic Stem Cell Self-Renewal to Specify Endoderm. CellReports 1–16.

Kalmar, T., Lim, C., Hayward, P., Muñoz-Descalzo, S., Nichols, J., Garcia-Ojalvo, J. and Arias, A. M. (2009). Regulated fluctuations in nanog expression mediate cell fate decisions in embryonic stem cells. PLoS Biol 7, e1000149.

Kang, M., Piliszek, A., Artus, J. and Hadjantonakis, A. K. (2012). FGF4 is required for lineage restriction and salt-and-pepper distribution of primitive endoderm factors but not their initial expression in the mouse. Development 140, 267–279.

Kunath, T., Saba-El-Leil, M. K., Almousailleakh, M., Wray, J., Meloche, S. and Smith, A. (2007). FGF stimulation of the Erk1/2 signalling cascade triggers transition of pluripotent embryonic stem cells from self-renewal to lineage commitment. Development 134, 2895–2902.

Lander, A. D., Gokoffski, K. K., Wan, F. Y. M., Nie, Q. and Calof, A. L. (2009). Cell lineages and the logic of proliferative control. PLoS Biol 7, e15–e15.

Marks, H., Kalkan, T., Menafra, R., Denissov, S., Jones, K., Hofemeister, H., Nichols, J., Kranz, A., Stewart, A. F., Smith, A., et al. (2012). The transcriptional and epigenomic foundations of ground state pluripotency. Cell 149, 590–604.

Morgani, S. M., Canham, M. A., Nichols, J., Sharov, A. A., Migueles, R. P., Ko,. S. H. and Brickman, J. M. (2013). Totipotent Embryonic Stem Cells Arise in Ground-State Culture Conditions. CellReports 3, 1945–1957.

Mulvey, C. M., Schröter, C., Gatto, L., Dikicioglu, D., Fidaner, I. B., Christoforou, A., Deery, M. J., Cho, L. T. Y., Niakan, K. K., Martinez Arias, et al. (2015). Dynamic Proteomic Profiling of Extra-Embryonic Endoderm Differentiation in Mouse Embryonic Stem Cells. STEM CELLS 33, 2712–2725.

Nichols, J., Silva, J., Roode, M. and Smith, A. (2009). Suppression of Erk signalling promotes ground state pluripotency in the mouse embryo. Development 136, 3215–3222.

Nichols, J., Zevnik, B., Anastassiadis, K., Niwa, H., Klewe-Nebenius, D., Chambers, I., Schöler, H. and Smith, A. (1998). Formation of pluripotent stem cells in the mammalian embryo depends on the POU transcription factor Oct4. Cell 95, 379–391.

Ohnishi, Y., Huber, W., Tsumura, A., Kang, M., Xenopoulos, P., Kurimoto, K., Oleś, A. K., Araúzo-Bravo, M. J., Saitou, M., Hadjantonakis, A.-K., et al. (2014) Cell-to-cell expression variability followed by signal reinforcement progressively segregates early mouse lineages. Nature Cell Biology 16, 27–37.

Plahte, E., Mestl, T. and Omholt, S. W. (1995). Feedback Loops, Stability and Multistationarity in Dynamical Systems. J. Biol. Syst. 03, 409–413.

Plusa, B., Piliszek, A., Frankenberg, S., Artus, J. and Hadjantonakis, A.-K. (2008). Distinct sequential cell behaviours direct primitive endoderm formation in the mouse blastocyst. Development 135, 3081–3091.

Ptashne, M. (2004). A Genetic Switch. CSHL Press.

Rossant, J. and Tam, P. P. L. (2009). Blastocyst lineage formation, early embryonic asymmetries and axis patterning in the mouse. Development 136, 701–713.

Schrode, N., Saiz, N., Di Talia, S. and Hadjantonakis, A.-K. (2014). GATA6 Levels Modulate Primitive EndodermCell Fate Choice and Timing in the Mouse Blastocyst. Dev Cell 1–14.

Shimosato, D., Shiki, M. and Niwa, H. (2007). Extra-embryonic endoderm cells derived from ES cells induced by GATA factors acquire the character of XEN cells. BMC Dev Biol 7, 80.

Singh, A. M., Hamazaki, T., Hankowski, K. E. and Terada, N. (2007). A heterogeneous expression pattern for Nanog in embryonic stem cells. STEM CELLS 25, 2534–2542.

Snoussi, E. H. (1998). Necessary Conditions for Multistationarity and Stable Periodicity. J. Biol. Syst. 06, 3–9.

Thomas, R. (1981). On the Relation Between the Logical Structure of Systems and Their Ability to Generate Multiple Steady States or Sustained Oscillations. In Springer Series in Synergetics (eds. Dora, Della, J., Demongeot, J., and Lacolle, B., pp. 180–193–193. Berlin, Heidelberg: Springer Berlin Heidelberg.

Wamaitha, S. E., del Valle, I., Cho, L. T. Y., Wei, Y., Fogarty, N. M. E., Blakeley, P., Sherwood, R. I., Ji, H. and Niakan, K. K. (2015). Gata6 potently initiates reprograming of pluripotent and differentiated cells to extraembryonic endoderm stem cells. Genes Dev 29, 1239–1255.

Xenopoulos, P., Kang, M., Puliafito, A., Di Talia, S. and Hadjantonakis, A.-K. (2015). Heterogeneities in Nanog Expression Drive Stable Commitment to Pluripotency in the Mouse Blastocyst. CellReports.

Yamanaka, Y., Lanner, F. and Rossant, J. (2010). FGF signal-dependent segregation of primitive endoderm and epiblast in the mouse blastocyst. Development 137, 715–724.

